# Anti-inflammatory properties of chemical probes in human whole blood: focus on prostaglandin E_2_ production

**DOI:** 10.1101/2019.12.30.890715

**Authors:** Filip Bergqvist, Yvonne Sundström, Mingmei Shang, Iva Gunnarsson, Ingrid Lundberg, Michael Sundström, Per-Johan Jakobsson, Louise Berg

## Abstract

We screened 57 chemical probes, high-quality tool compounds, and relevant clinically used drugs to investigate their effect on pro-inflammatory prostaglandin E_2_ (PGE_2_) production and interleukin-8 (IL-8) secretion in human whole blood. Freshly drawn blood from healthy volunteers and patients with systemic lupus erythematosus (SLE) or dermatomyositis was incubated with compounds at 0.1 or 1 μM and treated with lipopolysaccharide (LPS, 10 μg/mL) to induce a pro-inflammatory condition. Plasma was collected after 24 hours for lipid profiling using liquid chromatography tandem mass spectrometry (LC-MS/MS) and IL-8 quantification using enzyme-linked immunosorbent assay (ELISA). Each compound was tested in at least four donors at one concentration based on prior knowledge of binding affinities and *in vitro* activity. Our screening suggested that PD0325901 (MEK-1/2 inhibitor), trametinib (MEK-1/2 inhibitor), and selumetinib (MEK-1 inhibitor) decreased while tofacitinib (JAK inhibitor) increased PGE_2_ production. These findings were validated by concentration-response experiment in two donors. Moreover, the tested MEK inhibitors decreased thromboxane B_2_ (TXB_2_) production and IL-8 secretion. We also investigated the lysophophatidylcholine (LPC) profile in plasma from treated whole blood as these lipids are potentially important mediators in inflammation, and we did not observe any changes in LPC profiles. Collectively, we deployed a semi-high throughput and robust methodology to investigate anti-inflammatory properties of new chemical probes.

**Highlights:**

- Inhibitors for MEK decreased PGE_2_ and TXB_2_ production
- Inhibitors for MEK and ERK decreased IL-8 secretion
- JAK inhibitor tofacitinib increased PGE_2_ and TXB_2_ production

## Introduction

Inflammation is a highly controlled immune response to eliminate the cause of tissue injury or infection and to initiate tissue repair back to homeostasis via resolution [1, 2]. However, inflammation is not always terminated. Unresolved inflammation causes persistent pain, tissue degeneration, and loss of function. In particular, inflammatory responses drive many autoimmune diseases [3] and inflammation is a hallmark of cancer [4]. Thus, there is a great need for new therapies that are anti-inflammatory and safe.

Prostaglandin E_2_ (PGE_2_) is a potent lipid mediator of inflammation and immune responses, and PGE_2_ is a central mediator of pain, edema, and cartilage erosion typically observed in the joints of rheumatoid arthritis patients [5, 6]. In addition, PGE_2_ is a promotor of the immunosuppressive tumor microenvironment with major impact on tumor progression [4, 7, 8]. During inflammation, PGE_2_ is synthesized via conversion of arachidonic acid by cyclooxygenases (COX-1 and COX-2) into unstable PGH_2_ that is further metabolized by the inducible terminal synthase microsomal prostaglandin E synthase-1 (mPGES-1) to generate PGE_2_. Multiple non-steroidal anti-inflammatory drugs (NSAIDs) exist in clinical practice that unselectively decrease PGE_2_ production via inhibition of COX, but these drugs are all associated with adverse effects. Hence, selective inhibition of PGE_2_ production with small molecule inhibitors could therefore be a desirable therapeutic strategy in inflammation and cancer [9].

Interleukin-8 (IL-8) is a potent chemoattractant and activator of neutrophils. IL-8 signalling is implicated in multiple chronic inflammatory diseases [10] and cancer [11]. For example, a recent meta-analysis concluded that patients suffering from systemic lupus erythematosus (SLE) have increased levels of circulating IL-8 [12]. Patients with central neuropsychiatric SLE have increased concentration of IL-8 in cerebrospinal fluid compared to patients with non-central neuropsychiatric SLE [13]. IL-8 is also associated with renal damage and pulmonary fibrosis in SLE patients [14, 15]. Given that IL-8 is a stimulant for neutrophil activation, which plays a significant role in the pathogenesis of SLE [16], targeting IL-8 secretion or signalling could constitute a therapeutic strategy for SLE. A similar role of neutrophils and net formation has been reported in patients with dermatomyositis [17, 18]. In cancer, IL-8 is highly expressed in several types of cancer tissues [19] and serum concentration of IL-8 correlates with tumour burden [20]. The tumour-favouring actions of IL-8 include promotion of angiogenesis, increased survival of cancer stem cells, and attraction of myeloid cells that indorse the immunosuppressive tumour microenvironment [20].

In this study, we aimed to evaluate the effect of 57 chemical probes, high-quality tool compounds, and relevant control drugs on eicosanoid production and IL-8 secretion in human whole blood. A chemical probe is defined as “… a selective small-molecule modulator of a protein’s function that allows the user to ask mechanistic and phenotypic questions about its molecular target in biochemical, cell-based or animal studies” [21], and these compounds follow the criteria of *in vitro* potency (IC_50_ or Kd <100 nM), high selectivity versus other protein subfamilies (>30-fold), and on-target cell activity at 1 μM. The chemical probes and other high-quality tool compounds included are mainly epigenetic modulators and kinase inhibitors that were produced in academic collaborations or donated by pharmaceutical companies within the Structural Genomic Consortium (SGC, www.thesgc.org), which aims to investigate novel targets for drug development in open science and in collaboration with the pharmaceutical industry. These inhibitors were tested here at one concentration (in triplicates, n=4-15 donors) based on previous knowledge of binding affinities and toxicity *in vitro*, as assessed using other validated assays in our laboratories (https://ultra-dd.org/tissue-platforms/cell-assay-datasets).

## Materials and methods

### Ethical approval and consent to participate

Ethical approval for this study was granted by local research ethics committee at Karolinska University hospital (Dnr 02-196) and the Regional Ethical Review Board in Stockholm (Dnr 2015/2001-31/2). Full informed consent according to the Declaration of Helsinki was obtained from all patients.

### Collection of blood

Peripheral venous blood was drawn from 10 females and 6 males, aged between 27 and 81 years. Healthy controls (n=4) and two patient groups were included: systemic lupus erythematosus (n=9) and dermatomyositis (n=3). The blood was collected in tubes containing sodium heparin (1000 U/mL).

### Inhibitors

The inhibitors (chemical probes and other high-quality tool compounds) tested here were obtained through the SGC (www.thesgc.org) and supplied by different distributers (**Supplementary Table 1**). Inhibitors and control drugs (**Supplementary Table 1**) were reconstituted at 10 mM in DMSO (D2250, Sigma-Aldrich), aliquoted in Eppendorf tubes or 96-well plates, and kept at −80°C. A fresh aliquot was used at each experiment. Diclofenac (dual COX-1/2 inhibitor) was used as positive control for inhibition of prostanoid production. LPS (L6529, Sigma-Aldrich) was reconstituted in PBS (D8537, Sigma-Aldrich) to a final concentration of 0.1 mg/mL and kept at +8°C.

### Whole blood assay

Inhibitors and vehicle control (DMSO) were diluted in PBS at room temperature with no direct light on. The treatments were prepared in 25 μL portions to U-shaped 96-well plate and 200 μL of freshly drawn heparin blood (<2 hrs at room temperature) was added to the plate. The plate was incubated at 37°C for 30 min and then 25 μL of 0.1 mg/mL LPS in PBS was added followed by pipetting up and down 3 times (final concentration of LPS was 10 μg/mL). The tested concentration for inhibitor was 0.1 or 1 μM (Supplementary Table 1). The plate was incubated for 24 hrs at 37°C and then centrifuged at 3000 g for 10 min at 4°C. Working on ice, 100 μL plasma was recovered to a new plate (for prostanoid profiling) and from this 20 μL was transferred to a second plate (for IL-8 quantification). The plates were sealed with aluminum foil and stored at −80°C.

### Extraction of lipids

Plasma samples (80-240 μL) were thawed on ice and spiked with 50 μL deuterated internal standard mix containing 17 ng 6-keto-PGF_1α_-d4, 8 ng PGF_2α_-d4, 12 ng PGE_2_-d4, 8 ng PGD_2_-d4, 8 ng TXB_2_-d4, and 8 ng 15-deoxy-Δ12,14PGJ_2_-d4 (Cayman Chemical Company) prepared in 100% methanol. Protein precipitation was performed by addition of 800 μL 100% methanol, followed by vortexing, and centrifugation at 3000 g for 10 min at 4°C. The supernatants were collected in a new plate and evaporated under vacuum for 4 hrs. The evaporated samples (100-200 μL) were diluted to 1 mL with 0.05% formic acid in water and then loaded onto Oasis HLB 1cc 30mg plate (Waters Corporation, USA) that had been pre-conditioned with 1 mL of 100% methanol and 1 mL of 0.05% formic acid in water. The plate was washed with 10% methanol, 0.05% formic acid in water and lipids were eluted with 100% methanol. The eluates were dried under vacuum over-night and stored at −20°C until reconstituted in 50 μL of 20% acetonitrile in water prior to analysis with liquid chromatography tandem mass spectrometry (LC-MS/MS).

### Lipid profiling by LC-MS/MS

Lipids were quantified in negative mode with multiple reaction monitoring method, using a triple quadrupole mass spectrometer (Acquity TQ detector, Waters) equipped with an Acquity H-class UPLC (Waters). Eicosanoid were purchased from Cayman Chemicals and individually optimized for based on precursor ion m/z, cone voltage, collision energy, and fragment ion m/z (**Supplementary Table 2**). An eicosanoid mix containing all standards of interest was used to check interference in the LC-MS/MS analysis. LPC(14:0) and LPC(18:0) were used to set optimal analytical parameters for quantification of LPCs. Separation of lipids was performed on a 50 × 2.1 mm Acquity UPLC BEH C18 column 1.7 μm (Waters) with a 12 min stepwise linear gradient (20-95%) at a flowrate of 0.6 mL/min with 0.05% formic acid in acetonitrile as mobile phase B and 0.05% formic acid in water as mobile phase A. Data were analyzed using MassLynx software, version 4.1, with internal standard calibration and quantification to external standard curves for prostanoids. LPCs were normalized as area-% within each injection. Only lipids with peaks intensities of signal-to-noise greater than 10 (S/N >10) were considered in our data analysis.

### Quantification of IL-8

IL-8 was quantified in plasma by Human IL-8 (CXCL8) ELISA development kit (3560-1H, Mabtech) according to manufacturer’s instructions.

### Statistical analyses

Data are presented as mean±SEM if not stated otherwise. Statistical analyses were performed using GraphPad Prism 6 (GraphPad Sofware). One sample t-test was used to test significant difference. Statistical significance level was set to p<0.05.

## Results

### Development of whole blood assay

The whole blood assay was developed to screen for changes in multiple eicosanoids. Each eicosanoid and corresponding deuterated variant were individually optimized in the LC-MS/MS analysis. A dilution curve containing 6-keto PGF_1α_-d4, PGE_2_-d4, PGD_2_-d4, PGF_2α_-d4, TXB_2_-d4, 15d-PGJ_2_-d4, LTB_4_-d4, LTC_4_-d5, LTD_4_-d5, 5-HETE-d8, 12-HETE-d8, 15-HETE-d8, and undeuterated variants of 13-HODE, RvD1, RvD2, 17-hydroxy DHA, and protectin DX was spiked into 100 μL plasma at different stages throughout the extraction. A dilution curve was spiked in water at the same step. The dilution curve ranged from 0.006-1.5 pmol as final amount injected on the column in the LC-MS/MS analysis. This enabled us to investigate the lower limit of quantification (LLOQ), recovery efficacy, and matrix effect for each eicosanoid. The LLOQ injected on column was considered as great (0.02-0.05 pmol), good (0.1-0.2 pmol), or poor (0.4-1.5 pmol). Eicosanoids with great LLOQ were PGE_2_, PGF_2α_, TXB_2_, RvD1, RvD2, LTB_4_, protectin DX, and 13-HODE; good LLOQ were 6-keto PGF_1α_, PGD_2_, 5-HETE, 15-HETE, and LTD_4_; poor LLOQ were 15d-PGJ_2_, 12-HETE, 17-hydroxy DHA, and LTC_4_. The extraction recovery rates were 33-125%. The response in plasma compared to 20% acetonitrile were 52-116% due to matrix effects. The estimated LLOQ in 100 μL plasma was approximately 1 ng/mL for the best performing eicosanoids including PGE_2_, TXB_2_, PGF_2α_, RvD1, RvD2, and protectin DX. We can conclude that the method provided similar quantitative performance in plasma for many eicosanoids.

LPS at 0.1-10 μg/mL increased PGE_2_ and TXB_2_ production in human whole blood, which are the two dominant eicosanoids produced under these conditions [22]. All other eicosanoids were below the LLOQ. We chose 10 μg/mL of LPS as our final concentration, yielding a robust amount of PGE_2_ (49±4 ng/mL, n=5 donors) and TXB_2_ (24±9 ng/mL, n=5 donors). The prostanoid production was completely blocked using the dual COX-1/2 inhibitor diclofenac (10 μM). High concentration of DMSO (0.1%) slightly decreased PGE_2_ production by 20% (n=2 donors) while DMSO at 0.01% or 0.001% had no effect. The intra-assay coefficient of variation (CV, n=20 technical replicates) was 12% and 11% for PGE_2_ and TXB_2_, respectively. The inter-assay CV for control material (n=3 donors) was 20% for PGE_2_ and 30% for TXB_2_. This was performed on blood that was drawn, incubated, extracted, and analyzed at separate occasions. The suppression in signal due to matrix effects and/or recovery efficiency varied between donors and experiments, ranging from 10-70% suppression compared to signal in extracted blank (mean ± SD, n=6 donors, PGE_2_: 45±25%, TXB_2_: 40±20%). In summary, 24 hrs incubation of whole blood with 10 μg/mL LPS resulted in profound induction of the COX-1/2 products PGE_2_ and TXB_2_ that was efficiently blocked by diclofenac at 10 μM.

### Effect on PGE_2_ and TXB_2_ production

Our screening of inhibitors suggested that selected kinase inhibitors affected prostanoid production (**Figure 1**). The strongest reduction in PGE_2_ production was observed by MEK-1 inhibitor PD0325901 (31±6%, p=0.001, n=4) and MEK-1/2 inhibitor trametinib (34±7%, p<0.0001, n=15). Moderate suppression in PGE_2_ concentration was found for MEK-1/2 inhibitor selumetinib (65±9%, p=0.02, n=5), ERK-1/2 inhibitor SCH772984 (76±11%, p=0.04, n=13) and p38 inhibitor skepinone-L (76±8%, p=0.01, n=13). However, the tested p38 inhibitor pamapimod did not affect PGE_2_ production. Two of these compounds decreased TXB_2_ production, namely trametinib (63±6%, p=0.02, n=15) and selumetinib (74±7%, p=0.02, n=5). Diclofenac, here used as a positive control for inhibition of prostanoid production, blocked the prostanoid production while selective COX-2 inhibitor NS-398 inhibited only PGE_2_ production, in agreement with previously reported data for these compounds in whole blood assay [23]. The JAK inhibitor tofacitinib increased both PGE_2_ (286±51%, p=0.01, n=6) and TXB_2_ (169±20%, p=0.02, n=6) production. The IRAK-1/4 inhibitor I slightly increased the concentrations of PGE_2_ (139±15%, p=0.04, n=7) and TXB_2_ (133±8%, p=0.008, n=7).

**Figure 1.**
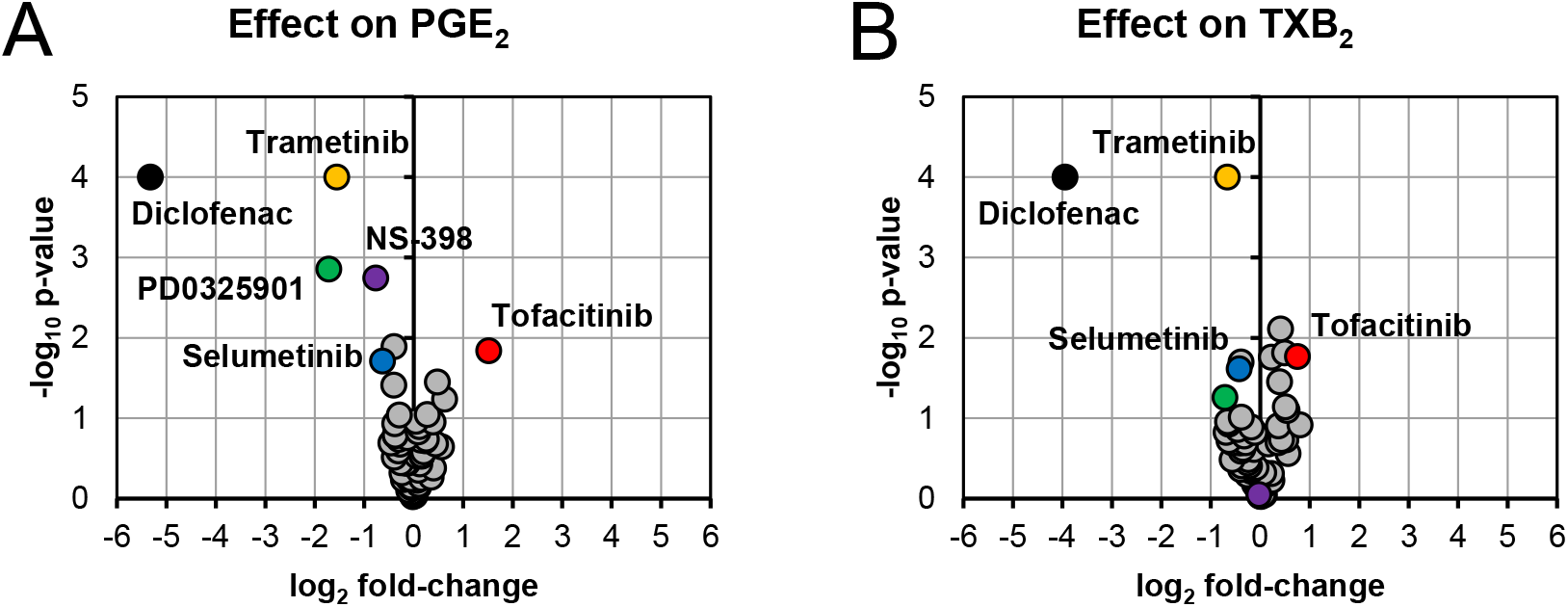
Volcano plots showing effects on PGE_2_ (A) and TXB_2_ (B) production in LPS-induced human whole blood. The top altered conditions compared to vehicle control based on fold-change (<0.5 or >2) and p-value (<0.05) are highlighted. Each inhibitor was tested in 4-15 donors. Statistical significance was tested using one sample t-test (p<0.05).

We chose to investigate the strongest observed effects in more detail by performing concentration-response experiments for PD0325901, trametinib, selumetinib, and tofacitinib. All three MEK inhibitors showed a concentration-dependent response on both PGE_2_ and TXB_2_ production while tofacitinib showed a concentration-dependent response on PGE_2_ production (**Figure 2**).

**Figure 2.**
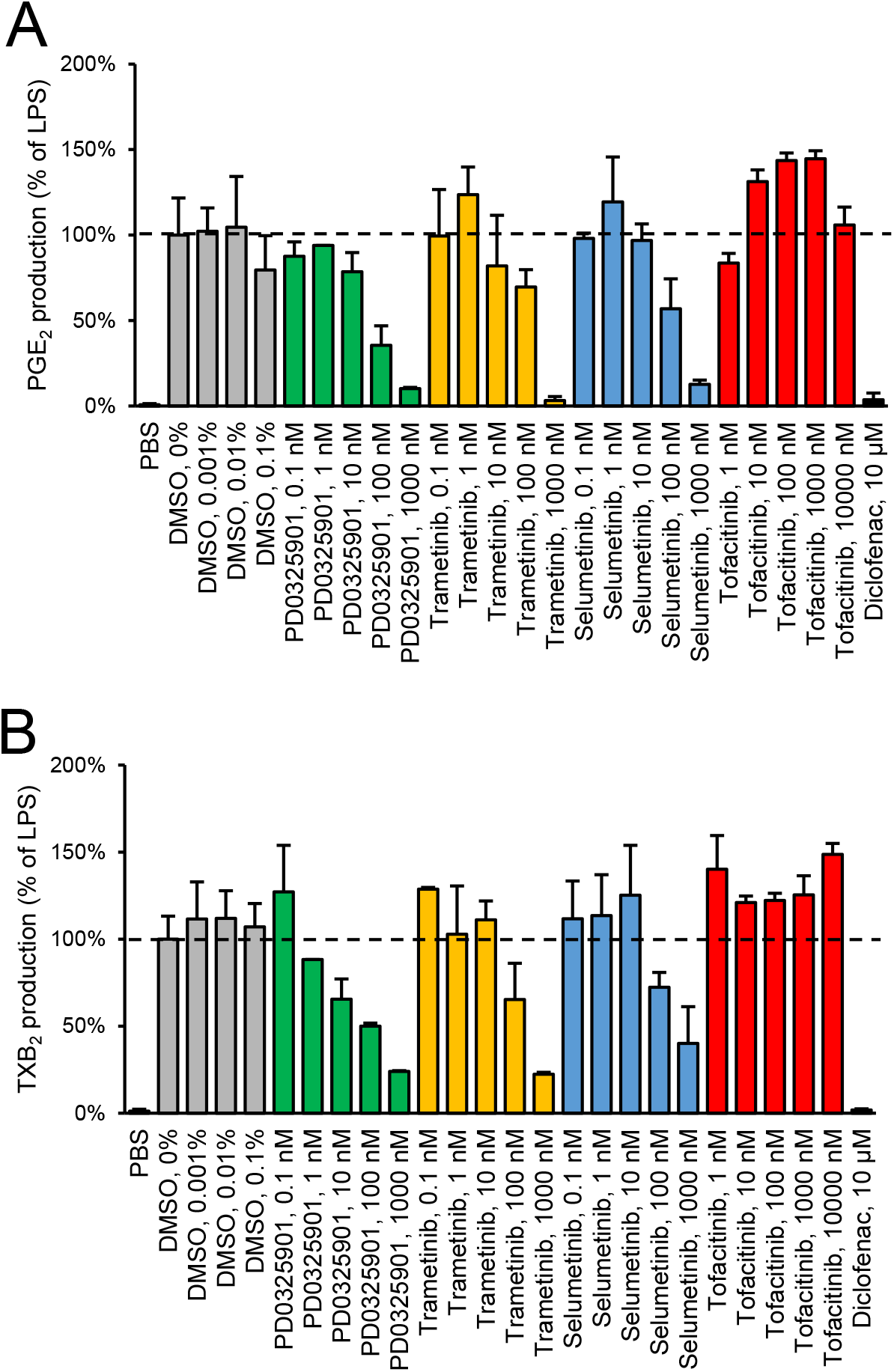
Validation of inhibitory effect on PGE_2_ production by MEK inhibitors in human whole blood. Diclofenac at 10 μM was used as positive control. Data are presented as mean±SD of biological replicates (n=2-6 per condition) from one representative experiment. The absolute PGE_2_ production in LPS control was 53.3±8.3 ng/mL. The concentration-response was tested in two donors.

### Effect on IL-8 secretion

In line with the effect on prostanoid production, reduction in IL-8 secretion was found for PD0325901 (24±9%, p=0.03, n=3), trametinib (27±5%, p<0.0001, n=13), and selumetinib (45±10%, p=0.03, n=3) (**Figure 3**). Moderate reduction in IL-8 secretion was found for SCH772984 (62±9%, p=0.002, n=12) and diclofenac (66±8%, p=0.003, n=11). We could also observe that tofacitinib increased IL-8 secretion (225±57%, p=0.16, n=3), however not with statistical significance.

**Figure 3.**
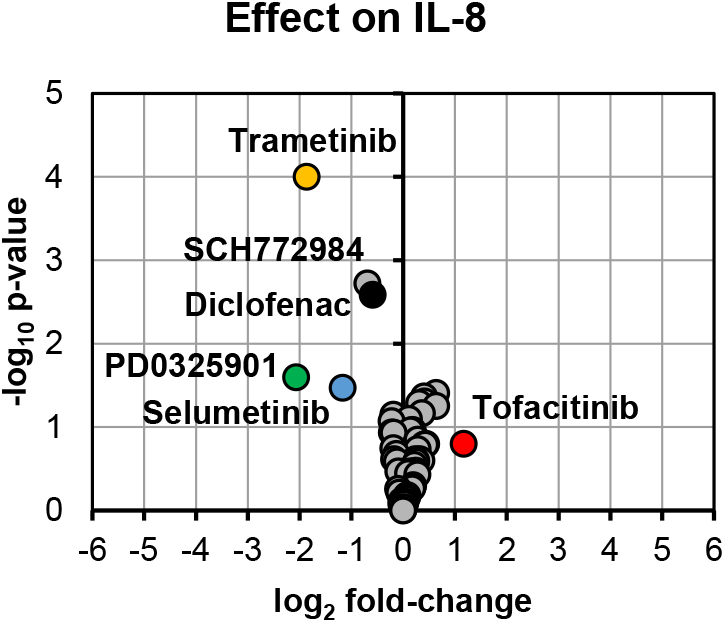
Volcano plot showing effects on IL-8 secretion in LPS-induced human whole blood. The top altered conditions compared to vehicle control based on fold-change (<0.5 or >2) and p-value (<0.05) are highlighted. Each inhibitor was tested in 3-13 donors. Statistical significance was tested using one sample t-test (p<0.05).

### Effect on LPC profile

We measured LPC species within our targeted LC-MS/MS analysis. LPCs are mainly generated by metabolism of membrane phosphatidylcholine by cytosolic phospholipase A_2_ [24]. These lipids have been reported to be involved in several cellular processes; sometimes with opposing effect depending on degree of saturation, concentration, and biological context [25, 26]. We observed no difference in total LPC or LPC profile when whole blood was treated with LPS neither did any of the tested inhibitors alter the LPC profile (**Figure 4**).

**Figure 4.**
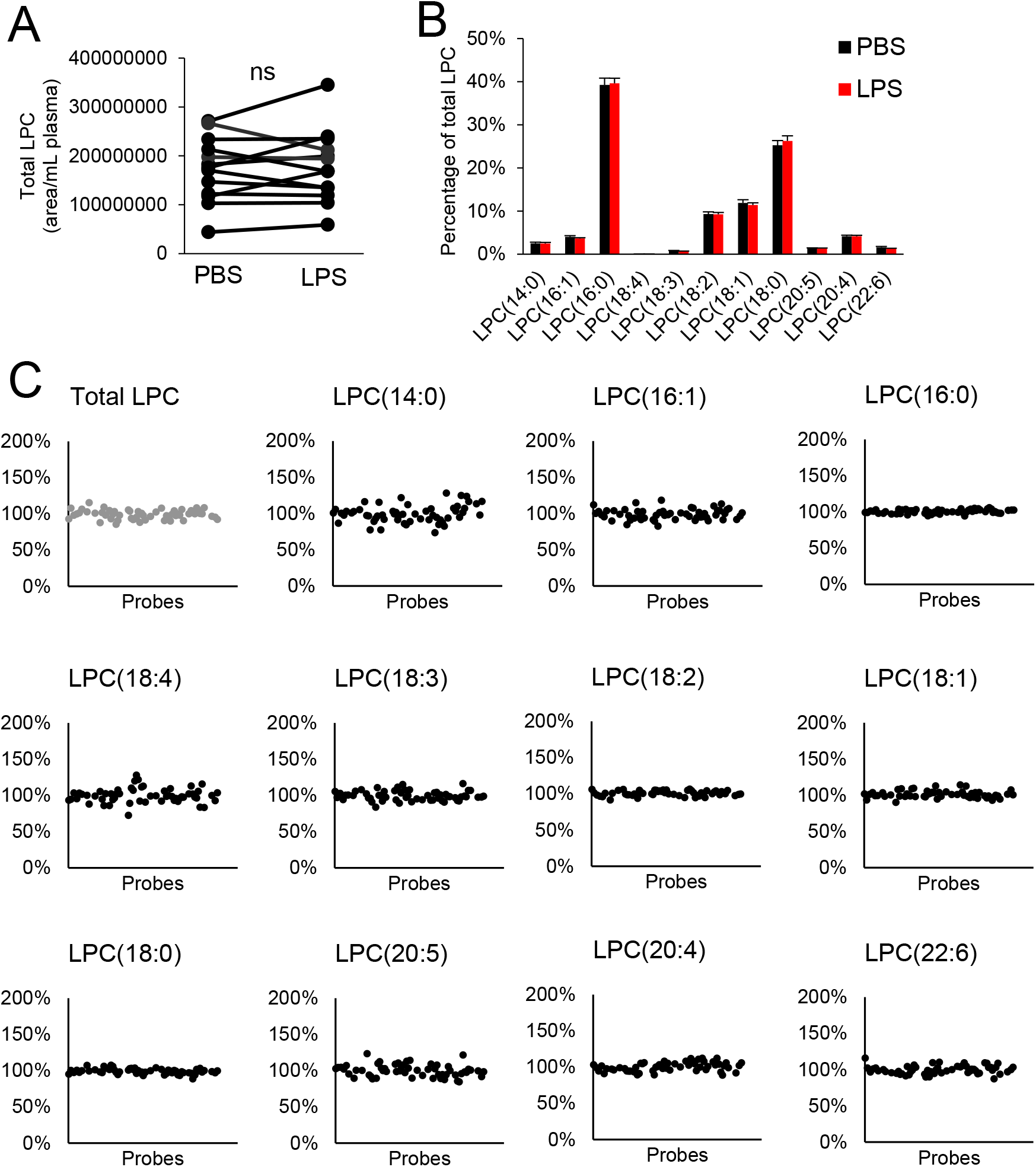
Effect on LPC profile in whole blood. There was no difference in total LPC (A) or LPC profile (B) with LPS treatment, and none of the tested compounds affected the LPC profile (C). Each inhibitor was tested in 4-15 donors.

## Discussion

We have tested the inhibitory effect on prostanoid production and IL-8 secretion in human whole blood for 57 high-quality inhibitors with known target specificities and *in vitro* potencies. None of the tested epigenetic modulators, which are acting on demethylases, bromodomains, or methyltransferases, affected PGE_2_ or IL-8 concentration. Inhibition of MEK-1/2 or ERK decreased PGE_2_ production and IL-8 secretion in this assay. This effect was observed for allosteric inhibitor trametinib (MEK-1/2), non ATP-competitive inhibitors PD0325901 (MEK-1) and selumetinib (MEK-1/2), and ATP-competitive inhibitor SCH772984 (ERK-1/2). These kinase targets are part of the RAS/RAF/MEK/ERK signaling transduction pathway, where inhibition of MEK prevents the downstream phosphorylation and activation of ERK that ultimately regulates cellular responses such as survival, lipid metabolism, and protein translation [27]. For example, MEK-1/2 inhibitor PD184352 decreased PGE_2_ production in melanoma cell line by decreased COX-2 expression due to inhibition of phosphorylation on ERK [28] and trametinib reduced IL-8 production in melanoma cell line [29]. We found that our positive control diclofenac for blocking prostanoid production decreased IL-8 secretion, which is explained by the fact that PGE_2_ stimulates IL-8 production in cultured cells [30–33]. While our study mainly focused on identifying inhibitory effects, we observed that JAK inhibitor tofacitinib increased both PGE_2_ production and IL-8 secretion. Tofacitinib is used to treat rheumatoid arthritis and it is known that tofacitinib can increase the expression of pro-inflammatory mediators, including PGE_2_, in macrophages by acting inhibitory on the expression of anti-inflammatory IL-10 [34]. The increased formation of pro-inflammatory PGE_2_ and platelet activating thromboxane A_2_ (as measured by stable metabolite TXB_2_) in human whole blood may be associated with the recently recognized increased risk of thromboembolism associated with JAK inhibitors in treatment of rheumatoid arthritis [35]. We acknowledge that the limitation of our study is the usage of one concentration per tested inhibitor. However, the used concentrations were based on reported IC_50_ and/or EC_50_ values as well as solid experiences in our laboratories using other validated assay systems (https://ultra-dd.org/index.php/tissue-platforms/cell-assay-datasets). We also demonstrated in concentration-response experiments that greater inhibitory effect could be achieved by increasing the concentration for the MEK inhibitors. However, this increases the risk of off-target effects and/or introduction of cellular toxicity that needs to be taken into account in experimental design and interpretation of results. In conclusion, we identified inhibitors for MEK or ERK as anti-inflammatory hits in our human whole blood assay. Based on the suppression in PGE_2_ production and IL-8 secretion, further investigation of the MEK/ERK signaling pathway may inform future therapeutic strategies to treat inflammatory diseases such as SLE and dermatomyositis.

## Acknowledgements

This work was supported by grants from Innovative Medicines Initiative (EU/EFPIA, ULTRA-DD, grant no: 115766), the Swedish Research Council (grant no: 2017-02577), Stockholm County Council (ALF, grant no: 20160378), The Swedish Rheumatism Association (grant no: R-755861), King Gustaf V’s 80 Years Foundation (grant no: n/a), and funds from Karolinska Institutet (grant no: n/a).

## Author contributions

F.B., Y.S., M.S., P-J.J., and L.B. contributed to study conception and design. F.B., Y.S., and M.M. performed experiments. F.B. analysed data, performed statistical analysis, and drafted the manuscript. I.G. and I.L. facilitated administrative, technical, or material support. All authors critically revised and approved the final version of the manuscript.

## Supplementary material

Supplementary Table S1.

Supplementary Table S2.

