## Supplementary Table 1 for "Anti-inflammatory properties of chemical probes in human whole blood: focus on prostaglandin E_2_ production"

| **#** | **Inhibitor** | **Item No. (Supplier)** | **Target** | **Target class** | **Assay conc.** (µM) | **PGE_2_, TXB_2_, LPCs** | **IL-8** |
| --- | --- | --- | --- | --- | --- | --- | --- |
| 1 | A-196 | 18317 (Cayman Chemical) | SUV420H1, SUV420H2 | Methyltransferase | 1 | n=4 | n=8 |
| 2 | A-366 | 16081 (Cayman Chemical) | EHMT2 (G9a), EHMT1 (GLP) | Methyltransferase | 1 | n=4 | n=8 |
| 3 | A-395 | SGC | EED (PRC2 complex) | Methyltransferase | 1 | n=5 | n=8 |
| 4 | BAY-598 | 18238 (Cayman Chemical) | SMYD2 | Methyltransferase | 1 | n=4 | n=8 |
| 5 | BAY-678 | SML1563 (Sigma-Aldrich) | Human neutrophil elastase | Serine protease | 1 | n=6 | n=13 |
| 6 | BAY-876 | SGC | GLUT1 | Glucose transporter | 1 | n=5 | n=5 |
| 7 | BAZ2-ICR | 17448 (Cayman Chemical) | BAZ2A, BAZ2B | Bromodomain | 1 | n=4 | n=8 |
| 8 | BI-9564 | SGC | BRD9, BRD7 | Bromodomain | 1 | n=4 | n=8 |
| 9 | Bromosporine | 14119 (Cayman Chemical) | BRD (pan-BRD) | Bromodomain | 1 | n=4 | n=8 |
| 10 | CGI1746 | S7051 (Selleckchem) | BTK | Kinase | 0.1 | n=7 | n=13 |
| 11 | Diclofenac | D6899 (Sigma-Aldrich) | COX-1, COX-2 | MAPEG | 10 | n=12 | n=11 |
| 12 | Fingolimod | FTY720 (Sigma-Aldrich) | S1P receptor | GPCR | 0.1 | n=7 | n=13 |
| 13 | GSK2606414 | S7307 (Selleckchem) | PERK | Kinase | 0.1 | n=7 | n=13 |
| 14 | GSK2801 | 14120 (Cayman Chemical) | BAZ2A, BAZ2B | Bromodomain | 1 | n=4 | n=8 |
| 15 | GSK343 | 14094 (Cayman Chemical) | EZH2 | Methyltransferase | 1 | n=5 | n=8 |
| 16 | GSK484 | 17488 (Cayman Chemical) | PAD4 | Arginine deiminases | 1 | n=4 | n=10 |
| 17 | GSK591 | 18354 (Cayman Chemical) | PRMT5 | Methyltransferase | 1 | n=4 | n=8 |
| 18 | GSK864 | SGC | Mutant IDH1 | Dehydrogenase | 1 | n=4 | n=5 |
| 19 | GSK-J4 | 12073 (Cayman Chemical) | JMJD3, UTX (prodrug of GSK-J1) | Demethylase | 1 | n=5 | n=8 |
| 20 | GSKLSD1 | 16439 (Cayman Chemical) | KDM1A (LSD1) | Demethylase | 1 | n=5 | n=8 |
| 21 | I-BRD9 | 17749 (Cayman Chemical) | BRD9 | Bromodomain | 1 | n=4 | n=8 |
| 22 | I-CBP112 | 14468 (Cayman Chemical) | CREBBP, EP300 | Bromodomain | 1 | n=4 | n=8 |
| 23 | IOX1 | 11573 (Cayman Chemical) | KDM (Pan-2-OG) | Demethylase | 1 | n=4 | n=8 |
| 24 | IOX2 | 11572 (Cayman Chemical) | PHD2 | Hydroxylases | 1 | n=4 | n=10 |
| 25 | IRAK-1/4 inhibitor I | I5409 (Sigma-Aldrich) | IRAK-1, IRAK4 | Kinase | 1 | n=7 | n=13 |
| 26 | JQ1 | 11187 (Cayman Chemical) | BET | Bromodomain | 0.1 | n=4 | n=8 |
| 27 | LP99 | 17661 (Cayman Chemical) | BRD7, BRD9 | Bromodomain | 1 | n=4 | n=8 |
| 28 | MK-2206 | S1078 (Selleckchem) | AKT1, AKT2, AKT3 | Kinase | 0.1 | n=6 | n=13 |
| 29 | MS023 | 18361 (Cayman Chemical) | Type I PRMTs | Methyltransferase | 1 | n=4 | n=8 |
| 30 | MS049 | 18348 (Cayman Chemical) | PRMT4 (CARM1), PRMT6 | Methyltransferase | 1 | n=4 | n=8 |
| 31 | NI-57 | 17662 (Cayman Chemical) | BRPF1, BRPF2, BRPF3 | Bromodomain | 1 | n=4 | n=8 |
| 32 | NS-398 | N194 (Sigma-Aldrich) | COX-2 | MAPEG | 0.1 | n=8 | n=8 |
| 33 | NVS-1 | 18316 (Cayman Chemical) | CECR2 | Bromodomain | 1 | n=4 | n=8 |
| 34 | NVS-PAK1-1 | SGC | PAK-1 | Kinase | 0.1 | n=8 | n=13 |
| 35 | OF-1 | 17124 (Cayman Chemical) | BRPF1, BRPF2, BRPF3 | Bromodomain | 1 | n=4 | n=8 |
| 36 | OICR-9429 | 16095 (Cayman Chemical) | WDR5 (MLL complex) | Methyltransferase | 1 | n=4 | n=8 |
| 37 | Pamapimod | 20971 (Cayman Chemical) | p38 | Kinase | 0.1 | n=4 | not tested |
| 38 | PD0325901 | 13034 (Cayman Chemical) | MEK-1 | Kinase | 0.1 | n=4 | n=3 |
| 39 | PFI-1 | 11155 (Cayman Chemical) | BET | Bromodomain | 1 | n=4 | n=8 |
| 40 | PFI-2 | 14678 (Cayman Chemical) | SETD7 | Methyltransferase | 1 | n=5 | n=8 |
| 41 | PFI-3 | 15267 (Cayman Chemical) | SMARCA2, SMARCA4, PBRM1 | Bromodomain | 1 | n=4 | n=8 |
| 42 | PFI-4 | 17663 (Cayman Chemical) | BRPF1B | Bromodomain | 1 | n=5 | n=8 |
| 43 | SCH772984 | S7101 (Selleckchem) | ERK1, ERK2 | Kinase | 0.1 | n=13 | n=12 |
| 44 | Selumetinib | S1008 (Selleckchem) | MEK-1, MEK-2 | Kinase | 0.1 | n=4 | n=3 |
| 45 | SGC0946 | 13967 (Cayman Chemical) | DOT1L | Methyltransferase | 1 | n=4 | n=7 |
| 46 | SGC707 | 17017 (Cayman Chemical) | PRMT3 | Methyltransferase | 1 | n=5 | n=8 |
| 47 | SGC-CBP30 | 14469 (Cayman Chemical) | CREBBP, EP300 | Bromodomain | 1 | n=4 | n=8 |
| 48 | SGX-523 | S1112 (Selleckchem) | MET | Kinase | 0.1 | n=6 | n=13 |
| 49 | Skepinone-L | S7214 (Selleckchem) | p38 | Kinase | 0.1 | n=13 | n=13 |
| 50 | T-26c | SGC | MMP-13 | Matrix metalloproteinase | 1 | n=5 | n=5 |
| 51 | Tofacitinib | PZ0017 (Sigma-Aldrich) | JAK | Kinase | 1 | n=6 | n=3 |
| 52 | TP-064 | SGC | CARM1 (PRMT4) | Methyltransferase | 1 | n=5 | n=5 |
| 53 | Trametinib (GSK1120212) | S2673 (Selleckchem) | MEK1, MEK2 | Kinase | 0.1 | n=15 | n=13 |
| 54 | UNC0638 | 10734 (Cayman Chemical) | EHMT2 (G9a), EHTM1 (GLP) | Methyltransferase | 1 | n=4 | n=8 |
| 55 | UNC0642 | SGC | EHMT2 (G9a), EHTM1 (GLP) | Methyltransferase | 1 | n=4 | n=8 |
| 56 | UNC1215 | 13968 (Cayman Chemical) | L3MBTL3 | Methyl-lysine readers | 1 | n=5 | n=8 |
| 57 | UNC1999 | 14621 (Cayman Chemical) | EZH2 | Methyltransferase | 1 | n=4 | n=8 |

**Supplementary Table S1.** SGC: Structural Genomic Consortium.
