## Supplementary Table 2 for "Anti-inflammatory properties of chemical probes in human whole blood: focus on prostaglandin E_2_ production"

| **Analyte** | **Item No.*** | **Retention time** (min) | **Precursor** (m/z) | **Fragment** (m/z) | **Cone voltage** (V) | **Collision energy** (V) |
| --- | --- | --- | --- | --- | --- | --- |
| 6-keto PGF_1α_ | 15210 | 3.2 | 369.1 | 245.2 | 53 | 21 |
| 6-keto PGF_1α_-d4 | 315210 | 3.2 | 373.1 | 249.2 | 53 | 21 |
| PGE_3_ | 14990 | 4.7 | 349.1 | 269.3 | 45 | 20 |
| TXB_2_ | 19030 | 5.0 | 369.2 | 169.2 | 30 | 20 |
| TXB_2_-d4 | 319030 | 5.0 | 373.2 | 173.2 | 30 | 20 |
| PGF_2α_ | 16010 | 5.7 | 353.3 | 309.3 | 45 | 23 |
| PGF_2α_-d4 | 316010 | 5.7 | 357.3 | 313.3 | 45 | 23 |
| PGE_2_ (1st ion) | 14010 | 6.0 | 351.1 | 271.3 | 38 | 19 |
| PGE_2_ (2nd ion) |  | 6.0 | 351.1 | 315.4 | 38 | 14 |
| PGE_2_-d4 | 314010 | 6.0 | 355.1 | 275.3 | 38 | 19 |
| PGD_2_ | 12020 | 6.2 | 351.1 | 271.3 | 38 | 19 |
| PGD_2_-d4 | 312020 | 6.2 | 355.1 | 275.3 | 38 | 19 |
| RvD2 | 10007279 | 6.4 | 375.5 | 175.2 | 35 | 30 |
| RvD1 | 10012554 | 7.0 | 375.5 | 141.0 | 30 | 20 |
| LTD_4_-d5 | 10006199 | 8.1 | 500.3 | 177.2 | 40 | 31 |
| LTC_4_-d5 | 10006198 | 8.2 | 629.4 | 272.2 | 50 | 31 |
| Protectin DX | 10008128 | 8.4 | 359.3 | 153.1 | 33 | 19 |
| LTB_4_ | 20110 | 8.5 | 335.3 | 195.2 | 40 | 17 |
| LTB_4_-d4 | 320110 | 8.5 | 339.3 | 197.2 | 40 | 17 |
| 15d-PGJ_2_ | 18570 | 8.8 | 315.2 | 271.2 | 17 | 24 |
| 15d-PGJ_2_-d4 | 318570 | 8.8 | 319.2 | 275.2 | 17 | 24 |
| 13-HODE | 38600 | 8.9 | 295.4 | 195.3 | 40 | 20 |
| 15-HETE | 34720 | 8.9 | 319.4 | 219.1 | 30 | 15 |
| 15-HETE-d8 | 334720 | 8.9 | 327.4 | 226.2 | 30 | 15 |
| 17(S)-HDoHE | 10009794 | 9.0 | 343.3 | 201.2 | 30 | 16 |
| 12-HETE | 34570 | 9.0 | 319.4 | 179.1 | 35 | 17 |
| 12-HETE-d8 | 334570 | 9.0 | 327.4 | 184.1 | 35 | 17 |
| 5-HETE | 34230 | 9.1 | 319.4 | 115.0 | 30 | 18 |
| 5-HETE-d8 | 334230 | 9.1 | 327.4 | 116.0 | 30 | 18 |

**Supplementary Table S2.** Experimental details for quantification of eicosanoids by LC-MS/MS. *Standards were purchased from Cayman Chemical Company.
